# Multitask Learning of Biophysically-Detailed Neuron Models

**DOI:** 10.1101/2023.12.05.570220

**Authors:** Jonas Verhellen, Kosio Beshkov, Sebastian Amundsen, Torbjørn V. Ness, Gaute T. Einevoll

## Abstract

The human brain operates at multiple levels, from molecules to circuits, and understanding these complex processes requires integrated research efforts. Simulating biophysically-detailed neuron models is a computationally expensive but effective method for studying local neural circuits. Recent innovations have shown that artificial neural networks (ANNs) can accurately predict the behaviour of these detailed models in terms of spikes, electrical potentials, and optical readouts. While these methods have the potential to accelerate large network simulations by several orders of magnitude compared to conventional differential equation based modelling, they currently only predict voltage outputs for the soma or a select few neuron compartments. Our novel approach, based on enhanced state-of-the-art architectures for multitask learning (MTL), allows for the simultaneous prediction of membrane potentials in each compartment of a neuron model, at a speed of up to two orders of magnitude faster than classical simulation methods. By predicting all membrane potentials together, our approach not only allows for comparison of model output with a wider range of experimental recordings (patch-electrode, voltage-sensitive dye imaging), it also provides the first stepping stone towards predicting local field potentials (LFPs), electroencephalogram (EEG) signals, and magnetoencephalography (MEG) signals from ANN-based simulations. It further presents a challenging benchmark for MTL architectures due to the large amount of data involved, the presence of correlations between neighbouring compartments, and the non-Gaussian distribution of membrane potentials.

## 1 Introduction

In the seven decades since Hodgkin and Huxley first described the action potential in terms of ion channel gating [25, 22, 21, 23, 24, 18, 6], the scientific community has gained a comprehensive understanding of how individual neurons process information. In contrast, however, the behaviour of large networks of neurons remains poorly understood. Experimental studies provide qualitative insights through statistical correlations between recorded neural activity and sensory stimulation or animal behaviour [26, 27, 29, 28], but statistical modelling offers little information on how networks perform neural computation or give rise to neural representations. Mechanistic modelling, in which detailed neuron models or networks of detailed neuron models are simulated on a computer, offers an alternative approach to studying the network dynamics of neural circuits [9, 11].

Thanks to recent pioneering efforts facilitated by large supercomputers, we are now able to construct simulations containing tens of thousands of model neurons that mimic specific cortical columns in mammalian sensory cortices [61, 55, 43, 58, 5]. Even more recent advances have reduced the need for supercomputers by distilling the output of biophysically-detailed neuron models into easier-to-evaluate artificial neural networks (ANN) [4, 51].

In the present paper we explore accelerated techniques for biophysically-detailed neuron models with full electrophysiological detail, a task that was previously deemed impractical due to the high computational cost involved. Our approach builds on previous work that focused on predicting outgoing action potentials or other experimental variables in a limited number of compartments. However, our approach is unique in that it allows for the simultaneous prediction of the membrane potentials (and membrane currents) for all compartments, see Figure 1. This allows for future comparison of model output with a wider range of experimental recordings of membrane potentials (dendritic patch electrodes, voltage-sensitive dye imaging), and also the calculation of extracellular signals such as local field potentials (LFPs), electroencephalogram (EEG) signals, and magnetoencephalography (MEG) signals [17]. This approach is an important step forward in our ability to simulate and eventually better understand the functioning of the brain in health and disease.

**Figure 1:**
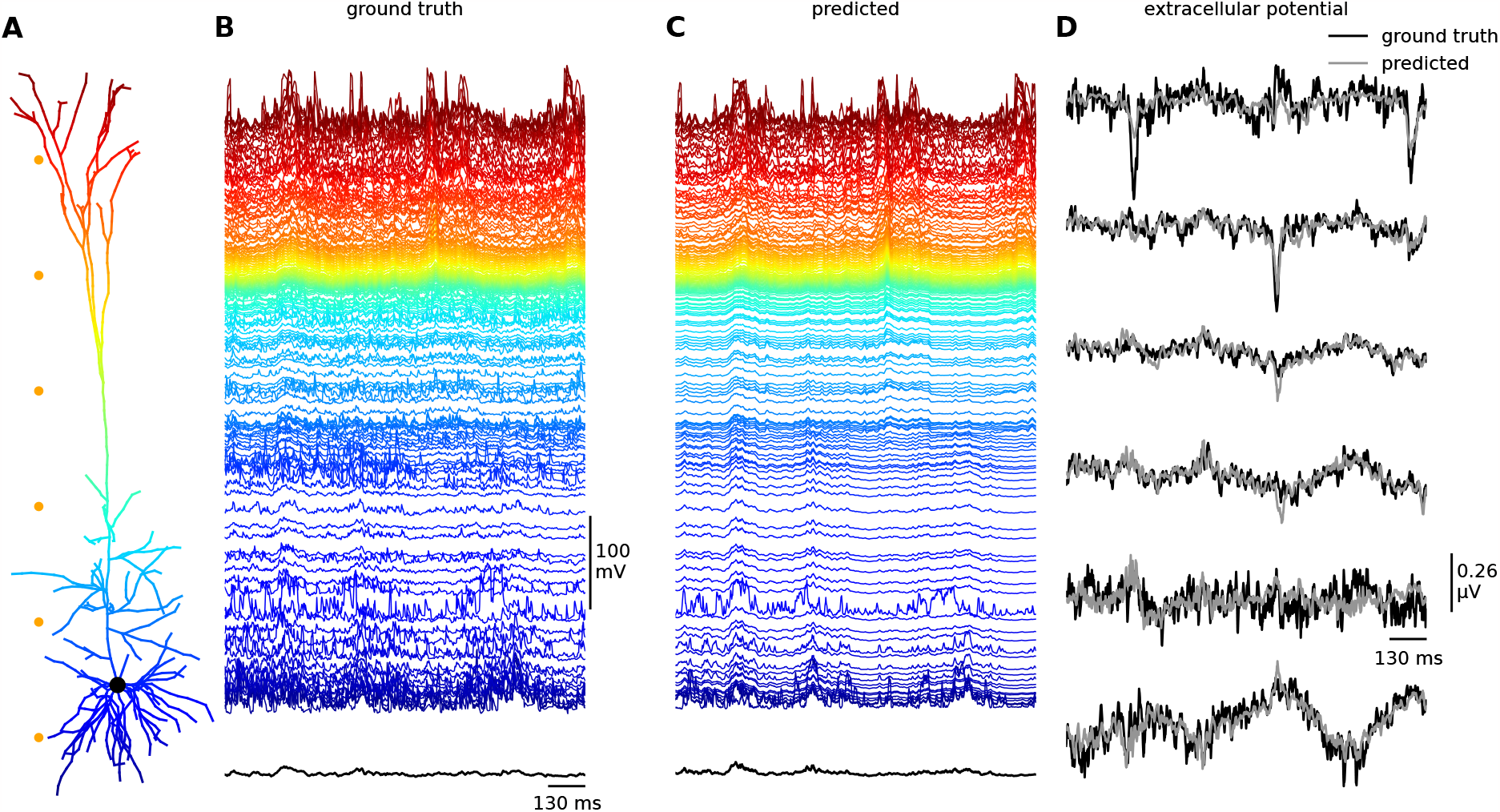
A) Illustration of the biophysically-detailed model with 639 compartments of a cortical layer V pyramidal cell model [19], which is the main object of study in this paper, color-coded by the compartment depth (consistent across all panels). B) Membrane voltages as calculated by a biophysically-detailed simulations of the multi-compartment model, used as the ground truth throughout this paper. C) Membrane voltages as predicted by our best-performing multi-task learning architecture, one time step at a time with a the time resolution of 1 ms for the prediction. D) Comparison between the ground truth and predicted extracellular potentials as calculated at six points outside the neuron (orange dots in panel A).

By predicting all membrane potentials across a biophysically-detailed neuron model simultaneously, rather than in the soma only as recently done by [4], we have made the transition from single-task learning to multi-task learning (MTL). In contrast to single-task learning that requires a separate model to be trained for each target, MTL optimises a single artificial neural architecture to predict multiple (heterogeneous) targets simultaneously. MTL approaches aim to improve generalisation and efficiency across tasks by leveraging statistical relationships between multiple targets [13, 62, 57, 34, 42].

In our work, statistical relationships between tasks, i.e. membrane potential predictions, arise from the correlations between compartments due to the biophysical mechanisms of the ion currents running through a neuron. To capture shared patterns that could be missed in single-task learning, multi-task learning generally relies on either one of two categories of neural architectures, respectively known as hard parameter [7] or soft parameter sharing models [10].

Hard parameter sharing models have a shared bottom layer in the neural network, while the output branches are task-specific. This means that the lower layers of the neural network are identical for all tasks, and only the final layers are tailored to each individual task. In contrast, soft parameter sharing models use dedicated sets of learning parameters and feature mixing mechanisms, allowing each task to have its own set of features and learning parameters while still efficiently sharing information. The choice between soft and hard parameter sharing models often depends on the nature of the tasks being learned. For example, if there is a substantial amount of overlap between the features necessary for each task, a hard parameter sharing model may be more appropriate, while if each (or any) task requires its own unique set of features for effective learning, a soft parameter sharing model may be more effective.

In this paper, we explore the capacity of various MTL architectures for distilling the full electrophysiology of a multi-compartment, biophysically-detailed layer 5 pyramidal neuron model [19]. Specifically, we compare a single type of hard parameter sharing model with two novel versions of state-of-the-art soft parameter sharing models, known as Multi-gate Mixture-of-Experts (MMoE) [41] and Multi-gate Mixture-of-Experts with Exclusivity (MMoEEx) [2]. Notably, in our computational experiments, we observed that the soft parameter sharing models substantially outperformed the hard parameter sharing model in the knowledge distillation of the electrophysiology of a biophysically-detailed neuron model. These findings highlight the potential benefits of soft-parameter sharing models for distilling complex electrophysiological data from biophysically-detailed neuron models, which we expect to have broad implications for computational neuroscience research.

For a comprehensive understanding of the MTL architectures utilised in this study, along with corresponding network diagrams, please refer to the detailed information provided in the Methods and Materials section. To make the following sections more accessible however, we briefly highlight that MMoE and MMoEEx, rely on a learnable feature mixing mechanism that controls the contribution of each learned data representation to the prediction of each task. The amount of learnable parameters in the feature mixing mechanisms grows linearly with the flattened input data size (batch size excluded). For the neural input data considered in this paper, the memory requirements regarding the amount of learnable parameters becomes prohibitively large. As a solution, we extend the MMoE and MMoEEx architectures to contain a compressor module that fixes the amount of learnable parameters in the feature mixing mechanisms to a predetermined number.

## 2 Results

To test the capability of different MTL architectures in accurately representing the full dynamic membrane potential of each of the 639 compartments of the large biophysically-detailed neuron model, we trained the models on a balanced dataset of simulated neural activity. For each target - a collection of membrane potentials (and the presence or absence of an outgoing action potential) - in the dataset, we provide a 100 ms history of neural activity and synaptic inputs from all compartments of the biophysically-detailed neuron model to the MTL models. Further details regarding the dataset and pre-processing steps can be found in the Methods and Materials section. To facilitate a fair comparison between MTL methods, we constructed a hard parameter sharing model (14 million trainable parameters) that is close in size to the soft parameter sharing models (12 million trainable parameters). Similarly, we used the same training procedures for each of the models – Adam [30] with or without task-balancing [38] – and evaluated the inference speed in a single session on publicly available hardware.

### 2.1 Multitask Prediction of Membrane Potential Dynamics in Biophysically-Detailed Neuron Models

After conducting comprehensive training on all three Multi-Task Learning (MTL) models, namely Hard Parameter Sharing (MH), Multi-gate Mixture-of-Experts (MMoE), and Multi-gate Mixture-of-Experts with Exclusivity (MMoEEx), we observed that the soft parameter sharing models exhibited a superior performance compared to the hard parameter sharing model, not only in terms of training but also in terms of generalization loss, see Figure 2 which showcases the training progress and the respective losses for each model. During the training process, we closely monitored the performance of each model through Tensorboard [1]. The hard parameter sharing model achieved a minimal training loss of 0.439 and a minimal validation loss of 0.680. The MMoE model, on the other hand, demonstrated better progress, reaching a minimal training loss of 0.241 and a minimal validation loss of 0.533. Similarly, the MMoEEx model exhibited better training results than the hard parameter sharing model, with a minimal recorded training loss of 0.262 and a minimal validation loss of 0.551.

**Figure 2:**
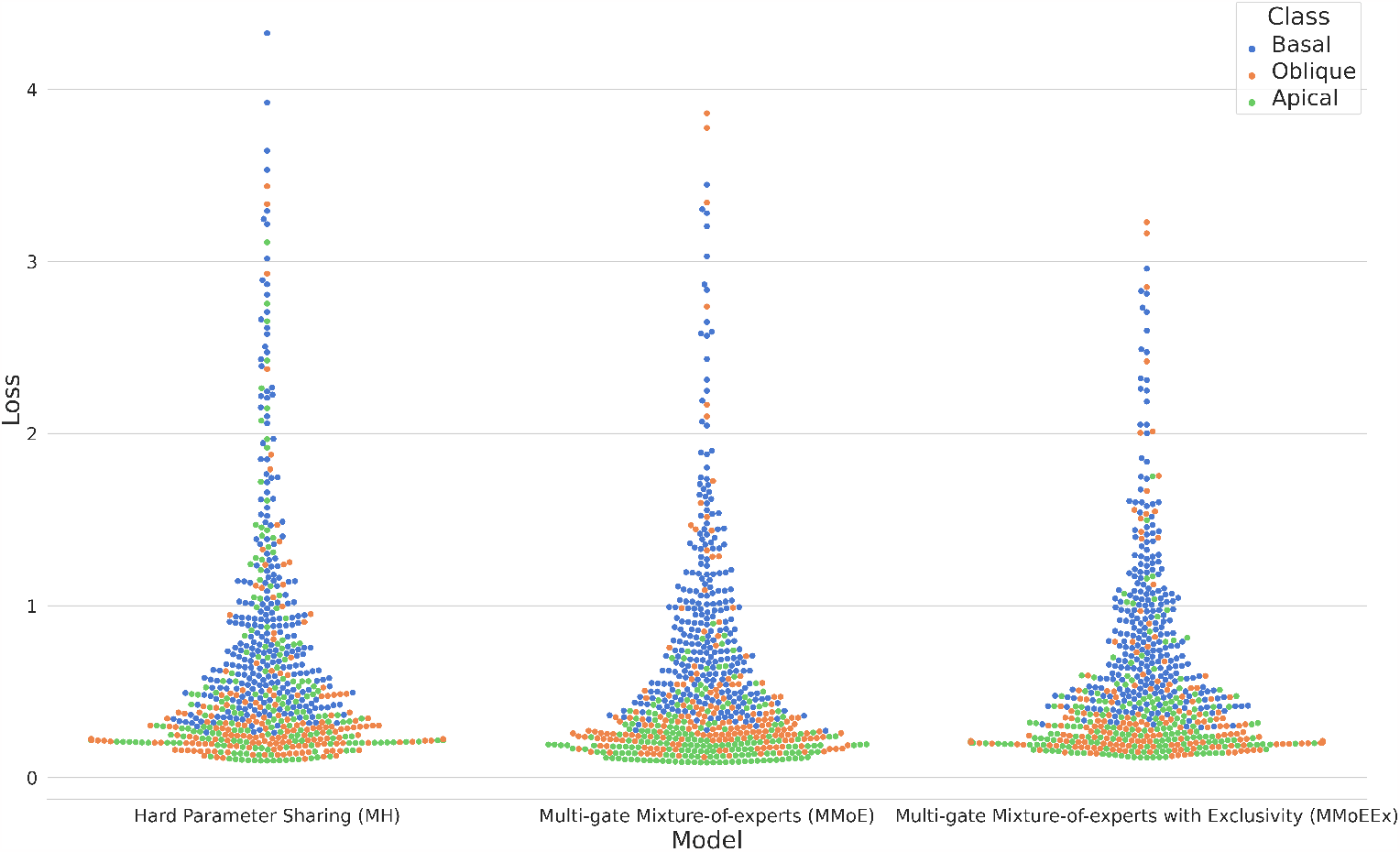
Swarm plots of the generalisation loss for each of the compartments (basal, oblique, apical) of the neuron model for each of the three models: Hard Parameter Sharing (MH), Multi-gate Mixture-of-experts (MMoE), and Multi-gate Mixture-of-Experts with Exclusivity (MMoEEx). Note that basal dendrites of a neuron receive incoming signals from other neurons and convey them towards the cell body, while the apical dendrite extends from the cell body to integrate signals from distant regions, and oblique dendrites play a role in the integration of synaptic inputs at various angles away from the other dendrites. The vertical axis represents the numeric loss values while the horizontal axis, the different ANN models are indicated. The density of the data points in a specific region indicates how many compartments have a similar loss value for each model.

For further evaluation, we focused on the MMoE model, as it showcased the best performance based on the validation results. By utilising the model weights saved at its optimal validation performance, we were able to predict the membrane potential of a compartment within the biophysically-detailed neuron model with a root mean squared error of 3.78 mV compared to a standard deviation of 13.20 mV across the compartments and batches of the validation data. As explained in more detail in the Discussion section, the electrophysiological data upon which the biophysically-detailed neuron model is built, has an experimental standard deviation of around 5 mV, depending on the type of neural activity. It is worth noting that the network models we explored primarily prioritised learning the membrane potentials of the neuron model’s compartments rather than accurately predicting spike generation in the soma. As a consequence, the presented results omit the performance analysis related to spike generation in the soma.

### 2.2 The Importance of Expert Diversity in MMoE and MMoEEx trained on Neural Data

Previous studies have suggested that higher diversity among experts in terms of the data representation they generate, see the Methods and Materials section, could potentially improve training and generalisation outcomes in MTL [41, 2], particularly for soft parameter sharing models. Motivated by these findings, we conducted a thorough investigation into the effect of expert diversity within the training process of the MMoE and MMoEEx models. To quantify and analyse expert diversity, we employed various diversity metrics. In addition to the diversity score, previously introduced by [2], which is based on the standardised distance matrix between experts, we extended our analysis to also include its determinant and permanent, see Methods. By considering these metrics, we were hoping to gain deeper insight into the level of diversity present among the experts throughout the training procedure. However, all three metrics exhibited the same temporal pattern. For a visual representation of these correlations and their implications on expert diversity, refer to Figure 1 in Appendix 1.

Interestingly, despite MMoEEx originally having been proposed in the literature as a means to enhance expert diversity, we made the intriguing observation that expert diversity exhibited significant variation across different training runs of the same model (MMoE or MMoEEx) with identical training data. This observation suggests that a soft parameter sharing model’s architecture alone does not guarantee any consistent trend in expert diversity, see the appendices for more details, and that multiple training runs are necessary for complex deep learning models to learn about their properties. Furthermore, we studied the strength of the expert weights used to predict each compartment in both MMoE and MMoEEx, see Figure 3. While larger average weight does not necessarily imply a better prediction, it corresponds to a larger contribution to the prediction coming from a particular expert. As one can see in both models, different experts focus on different parts of the neuron, with relatively higher mean weights in higher apical and basal regions. One noticeable difference is that in MMoEEx, the oblique parts of the neuron are more strongly represented, whereas somatic compartments are visibly different from zero in one of the experts.

**Figure 3:**
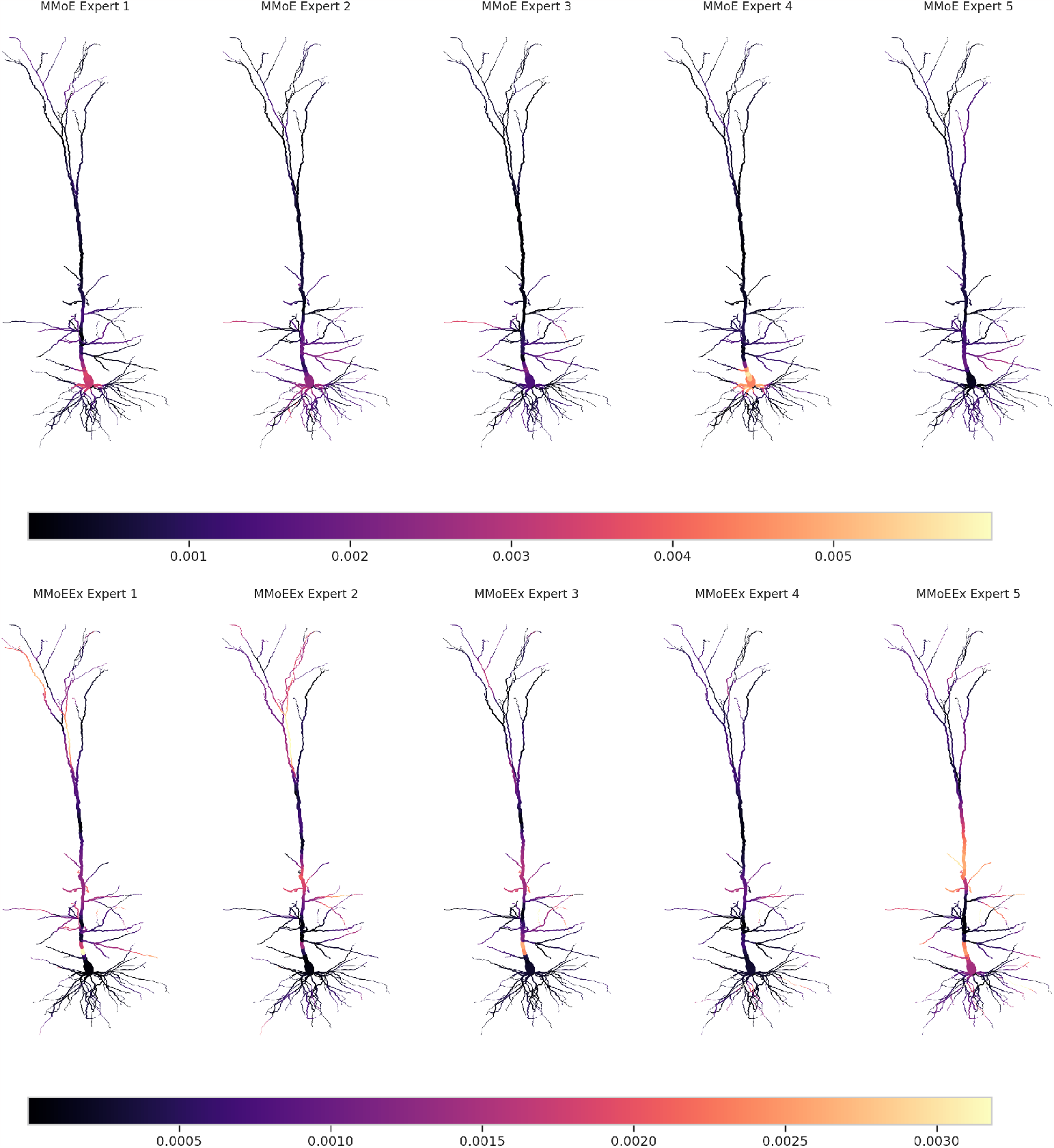
Mean weights from the experts in both Multi-gate Mixture-of-experts (MMoE; top row) and Multi-gate Mixture-of-experts with Exclusivity (MMoEEx; bottom row) after training the models on neural data, projected onto the different compartments of the cortical layer V biophysically-detailed neuron model. Each row consists of five subplots, representing different experts within the models. The color scale (normalised to the mean weights of the first expert) is indicated by the horizontal color bar located below each row.

### 2.3 Fast Simulation of Full Membrane Potential Dynamics of Multiple Neurons

Traditional simulation environments such as NEURON rely on numerical integration of compartment-specific differential equations that represent the active and passive biophysical mechanisms of the modeled neuron. This approach requires a significant amount of computational resources. One of the main attractive features of distilling biophysically-detailed neuron models into ANNs is the significant speed-up deep learning architectures can provide. Previous work from the literature, where only the output of the soma compartment was predicted, has shown that NEURON simulations of biophysically-detailed neuron model networks, consisting of up to 5000 model neuron instances, are up to five orders of magnitude slower than their ANN counterparts [51]. Note that, in network simulations, accelerators like GPUs can be used to compute independent timesteps - for different instances of the same model neuron - of the ANN model in parallel. In practice, the batch-size of the MTL models during inference can directly be interpreted as the number of model neurons being evaluated at the same time.

We recorded the mean and standard deviation runtimes of seven independent predictions for each of the three MTL models for different amounts of neurons and simulation times. In Table 1, we present the results of 100 neurons simulated for 1 ms, 10 ms and 100 ms, and for a 100ms simulation of 1000 neurons. To provide context, we also recorded the mean and standard deviation runtimes of seven independent 100 ms simulations of a single neuron in NEURON. Note that in the NEURON simulation, each timestep is passed to the next internally, while for the runtime tests of the deep learning models we are limited to running each timestep from Python causing an overhead for the MTL models. Clearly, predictions by MTL models are significantly faster than single-core simulations in NEURON. For the set-up with 1000 neurons, the hard parameter sharing model reaches an acceleration of 2 orders of magnitude over the NEURON simulation. Being able to predict voltages in parallel has a profound effect on efficiency, as a 1ms simulation of 100 neurons can be predicted in 0.45 seconds which is nearly half the time necessary for NEURON to simulate a single neuron for 100 ms, while requiring the same amount of update steps.

**Table 1:** Inference speed of all three MTL architectures - hard parameter sharing (MH), Multi-gate Mixture-of-experts (MMoE), and Multi-gate Mixture-of-experts with Exclusivity (MMoEEx) - after training on neural data generated by a biophysically-detailed model of a cortical layer 5 pyramidal neuron model compared to the classical NEURON simulation.

| Runtime (s) | Std. Dev. (ms) | Model | Hardware | Simulation (ms) | Batch Size |
| --- | --- | --- | --- | --- | --- |
| 1.01 | 29.7 | Neuron | CPU | 100 | 1 |
| 0.07 | 0.85 | MH | GPU | 1 | 100 |
| 0.45 | 1.03 | MMoE | GPU | 1 | 100 |
| 0.44 | 1.43 | MMoEEx | GPU | 1 | 100 |
| 0.70 | 25.4 | MH | GPU | 10 | 100 |
| 4.36 | 35.7 | MMoE | GPU | 10 | 100 |
| 4.35 | 15.0 | MMoEEx | GPU | 10 | 100 |
| 7.19 | 145 | MH | GPU | 100 | 100 |
| 43.6 | 58.6 | MMoE | GPU | 100 | 100 |
| 43.5 | 82.4 | MMoEEx | GPU | 100 | 100 |
| 10.9 | 153 | MH | GPU | 100 | 1000 |
| 365.0 | 34.2 | MMoE | GPU | 100 | 1000 |
| 365.0 | 33.5 | MMoEEx | GPU | 100 | 1000 |

Also note that while prediction times for the MTL models scale roughly linearly with simulation times, they do not scale linearly with batch size. Within the maximum batch size that can be accommodated by the GPU memory, the number of simulated neurons should only affect the evaluation time of a neural network through the previously discussed Python overhead, as is seen in the recorded prediction times for the hard parameter sharing model. However, for the soft parameter sharing models which contain matrix multiplications in the gates we observe a steep increase in runtime for large batch sizes. In general, it is worth remarking that MMoE and MMoEEx demonstrate inference runtimes which are significantly slower compared to the hard parameter sharing model. This discrepancy can be attributed to the hard parameter sharing model having a lower number of convolution filters and the soft parameter sharing models containing large and computationally expensive matrix multiplications.

## 3 Discussion

### 3.1 Multitask Prediction of Membrane Potential Dynamics in Biophysically-Detailed Neuron Models

In our investigation, we evaluated three advanced multi-task learning (MTL) neural network architectures [7, 41, 2], including MMoE and MMoEx which were augmented with a novel compressor module, to tackle the challenging task of capturing intricate membrane potential dynamics in a complex, multi-compartment, biophysically-detailed model of a layer 5 pyramidal neuron [19]. Given the considerable complexity of the model comprising 639 compartments, each generating distinct timeseries data potentially containing spikes, this MTL problem presented an exceptionally difficult distillation task. Nevertheless, our findings are encouraging, with the MMoE model achieving the lowest validation loss and demonstrating a root mean squared error of 3.78 mV in predicting membrane potential across all compartments. In comparison, the root mean squared error for the MMoEEx and MH models was 3.90 mV and 4.81 mV respectively. Notably, these errors are considerably smaller than both the standard deviation of our test set (13.20 mV) and the experimentally measured standard deviations of the peak membrane voltages during perisomatic step current firing (4.97 mV - 6.93 mV) or back-propagating action potential Ca^2+^ firing (5 mV) in layer 5 pyramidal neurons [33, 19].

Experimental recordings of the after-hyperpolarization depth of the membrane potential in the soma of layer 5 pyramidal neurons have a slightly lower standard deviation (3.58 mV - 5.82 mV) than electrophysiological measurements of action potentials [31, 32, 19], indicating that a more sensitive evaluation of distilled neuron models could be based on their performance in specific neuronal scenario’s. Notably, thus far, all trained models did not manage to learn the binary somatic spike prediction task, a challenge that could potentially be mitigated through a computationally intensive hyperparameter search for *γ* (explained in the Methods and Materials section) to assign higher importance to this specific task during the learning process. Although task balancing methods for soft parameter sharing models were explored in accordance with the procedures detailed in the Methods and Materials section, our findings indicate no improvement in performance, as outlined in Appendix 2. Furthermore, it is essential to highlight that these MTL models are anticipated to find utility in accelerating LFP calculations, where the significance of subthreshold components in membrane potential dynamics is well-established [52, 54, 35]. For further insights into calculating extracellular potentials based on the deep learning model’s membrane potential predictions and an example of such a downstream prediction, please refer to Appendix 3.

### 3.2 Measuring Experts Diversity in MMoE and MMoEEx Trained on Neural Data

To test the conjecture that in soft parameter-sharing MTL models, high diversity between experts can be beneficial in training and generalisation, we computed several diversity measures. Indeed, initially it looked like the model with highest diversity (MMoE) performed the best, despite the fact that unlike its counterpart (MMoEEx), it has no explicit inductive bias towards expert diversity. However, a subsequent retraining of the models showed that the known diversity metrics are neither robust nor correlated with training or generalisation performance. Further exploration of the exclusivity hyperparameter *α* might lead to more desirable results for the MMoEEx model. Additionally, increasing the number of experts should only lead to improvements in training and generalisation if a substantial amount of independence between the tasks had not yet been incorporated. Future work can make these statements more precise by studying the contribution of individual experts to the prediction of membrane potentials in morphologically distinct neuronal compartments.

### 3.3 Fast Simulation of Full Membrane Potential Dynamics of Multiple Neurons

Biophysically-detailed neuron models distilled into ANNs can be evaluated at significantly higher speeds than their classical counterparts. Because a single instance of a deep learning model can be used to predict outputs for multiple instances of the same neuron model in parallel, on accelerators such as GPUs, these models are particularly suited to accelerating large networks of model neurons. Previous results from the literature [51] have shown that using deep learning models that only predict the output voltage of the soma, could result in a five order magnitude speed-up for a network model of 5000 neurons. In this paper, we have shown that a speed-up of two orders of magnitude can be obtained for a network of 1000 neurons by making use of MTL deep learning models, when the voltage traces for each compartment of the underlying biophysically-detailed neuron model need to be predicted. The highest speedup was achieved with the MH model, which has a slightly higher root mean squared error (4.81 mV) compared to the other two models, but still performs within experimentally observed variability [19].

The presented MTL models did not succeed in the binary somatic spike prediction task. As such, the models are not particularly suited for running recurrently connected neural network simulations. They can, however, be used for investigating, for example, LFP, EEG, or MEG signals from different types of predetermined synaptic input to neural populations, which has commonly been used in the literature to study the origin and information content of these brain signals [53, 36, 37, 59, 40, 45, 14, 48, 15, 39, 47, 60, 50, 44, 16]. Furthermore, in cases where the synaptic input is predetermined, the individual timesteps can be treated independently and in parallel. For example, if we assume a batch size of 1000, this can either correspond to simultaneously simulating 1000 timesteps of a single neuron, or one timestep for a population of 1000 neurons.

An important application of the biophysically-detailed neuron model is the calculation of LFPs and downstream EEG and MEG signals. Recent work has shown the importance of LFP recordings in validating large computational models of brain tissue [56], such as the mouse V1 cortical area, developed by the Allen Institute [5]. The Allen Institute mouse V1 model contains 114 distinct model neuron types, and could be accelerated (by several orders of magnitude) by 114 separate deep learning models each representing one such neuron type. The current biophysically-detailed Allen V1 model makes use of neuron models with passive dendrites which should be significantly easier to distil into an MTL architecture than the layer 5 pyramidal neuron model represented here. Further acceleration of the MTL models discussed in this paper could be achieved by running model inference on multiple GPUs or more advanced accelerators such as TPUs and IPUs.

## 4 Methods and Materials

### 4.1 Multicompartmental NEURON Simulations and Data Balancing

As a baseline for training and testing, we used an existing dataset of electrophysiological data generated in NEURON [4] based on a well-known biophysical-detailed and multi-compartment model of cortical layer V pyramidal cells [19]. This model contains a wide range of dendritic (Ca^2+^-driven) and perisomatic (Na^+^ and K^+^-driven) active properties which are represented by ten key active ionic currents that are unevenly distributed over different dendritic compartments. The data was generated in response to presynaptic spike trains sampled from a Poisson process. For the purposes of this paper, it is important to note that the biophysically-detailed model contains 639 compartments and 1278 synapses and that 128 simulations of the complete model for 6 seconds of biological time each were included before data balancing.

The subthreshold dynamics of the membrane potential in a compartment have small variations and as a result one would expect them to be easy to predict. However, suprathreshold deviations generated by action potentials propagating through the dendrites can be more problematic to predict. To address this issue, we implemented a form of data balancing. We first identified the time points at which somatic spikes occurred and afterwards we used them to create a dataset in which one third of the targets included a spiking event. Additionally, we standardised the membrane potential through z-scoring. We used input data consisting of membrane potentials and incoming synaptic events across all compartments during a 100 ms time window, and target data consisting of 1 ms of membrane potentials and a binary value for the presence or absence of a somatic spike.

### 4.2 Hard and Soft Parameter Sharing Architectures for Multitask Learning

We implemented a temporal convolutional network (TCN) architecture [3] to learn spatiotemporal relationships between inputs (synaptic events and membrane potentials) and outputs (membrane potentials), for both hard and soft parameter-sharing architectures for multi-task learning. While recurrent neural networks [20, 8] such as GRU’s and LSTM’s are commonly used for sequence tasks, research has shown that convolutional architectures can perform just as well or better on tasks like audio synthesis, language modelling, and machine translation [3]. A TCN is a one-dimensional convolutional network with a causal structure, which guarantees causality layer by layer through the use of causal convolutions, padding and dilation. TCNs preserve the temporal ordering of the input data, making it impossible for the network to use information from the future to make predictions about the past. This is important because it ensures that the network can be applied in real-world scenarios where only past information is available.

In early studies of modern multi-task learning, hard parameter sharing was used to share the initial layers (“the bottom”) and task-specific top layers (“the towers”) of the neural network architecture. This approach has the advantage of being scalable with increasing numbers of tasks, but it can result in a biased shared representation that favours tasks with dominant loss signals. To address this issue, soft parameter sharing architectures have been developed, which utilise dedicated representations for each task. In this study, we employ two soft parameter sharing models, the multi-gate mixture-of-experts (MMoE) [41] and the multi-gate mixture-of-experts with exclusivity (MMoEEx) [2]. These models combine representations learned by multiple shared bottoms (“the experts”) through gating functions that apply linear combinations using learnable and data-dependent weights. MMoEEx is an extension of MMoE that encourages diversity among expert representations by randomly setting a subsection of weights to zero.

In our study, we employed a three-layered TCN with varying channel sizes of 32, 16, and 8 and a kernel size of 10. We also implemented a dropout of 0.2 for both the bottom architecture (hard parameter sharing) and the expert architectures (soft parameter sharing). For each tower, we used a three-layered feed-forward network with ELU activation functions and 10 or 25 hidden nodes, depending on whether it was a soft or hard parameter sharing model, respectively. To ensure increased stability, we replaced the final RELU activation function in the original TCN implementation with a sigmoid activation function. In addition, we included an extra TCN expert as a compressor module in the soft parameter sharing models to reduce the size of the data before feeding it into the gating functions, see Figure 4, which helped to avoid quadratic growth of the trainable parameters with respect to the input data size. To improve efficiency, we implemented mixed precision and multi-GPU training for all three models. Our models were trained on up to six parallel RTX2080Ti GPUs, and we performed inference experiments on publicly available Tesla P100 GPUs via Kaggle.

**Figure 4:**
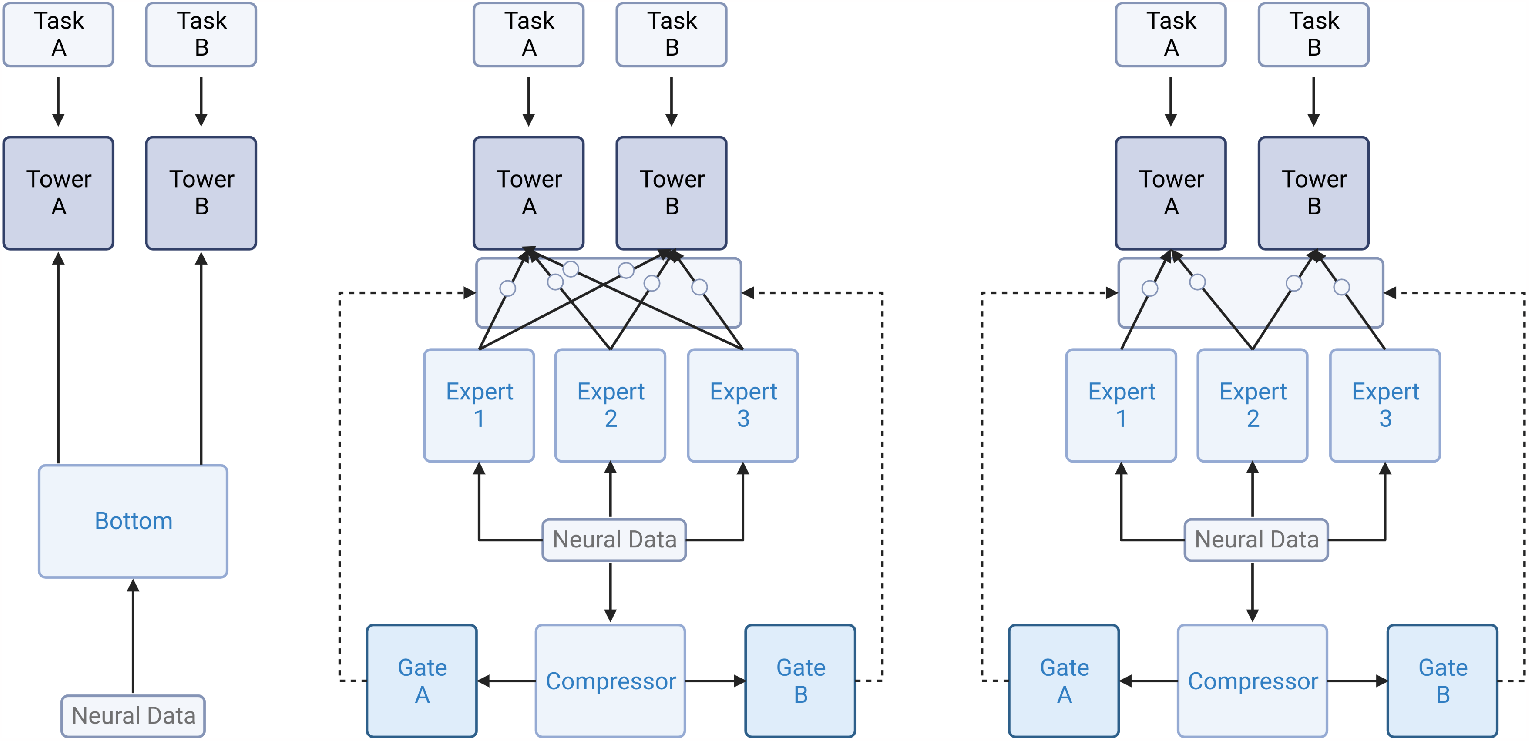
Schematic representation of the architectures of the hard parameter model (left), the MMoE model (middle), and the MMoEEx model (right).

### 4.3 Task Balancing and Expert Diversity

Effective multi-task learning sometimes requires some form of task balancing to reduce negative transfer or to prevent one or more tasks from dominating the optimisation procedure. To avoid these issues we make use of loss-balanced task weighting (LBTW) [38] which dynamically updates task weights in the loss function during training. For each batch, LBTW calculates loss weights based on the ratio between the current loss and the initial loss for each task, and a hyperparameter *α*. As *α* goes to 0, LBTW approaches standard multitask learning training. All taken together, the loss function can be summarised as

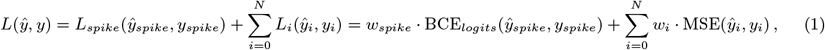

where *ŷ* and *y* are the target and the predicted data respectively, *w*_*spike*_ and *w*_*i*_ are the task weight, BCE_*logits*_ is the binary cross entropy loss combined with a sigmoid activation function, and MSE is the mean square error. The summation index i runs over all N compartments of the biophysically-detailed neuron model, and the LBTW task weights are recalculated every epoch *E*, for each batch *B*, according to

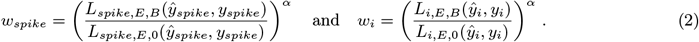

To measure the diversity in expert representations in MMoE and MMoEEx, we made use of diversity measurement proposed in the original MMoEEx paper. In that paper, the diversity between two experts *n* and *m* is calculated as a (real-valued) distance *d*(*n, m*) between the learned representations *f*_*n*_ and *f*_*m*_, as defined by

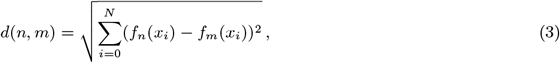

where *N* is the number of samples *x*_*i*_ in the validation set, which is used to probe the diversity of the experts. The diversity matrix *D* of a trained MMoE or MMoEEx model is defined by calculating all pairwise distances between expert representations as described above and normalising the matrix. In this normalised matrix, a pair of experts with distance close to 0 are considered near-identical, and experts with a distance close to 1 are considered to be highly diverse. To compare two different models in terms of overall diversity, we respectively define the first, second, and third diversity scores of a model as the mean entry 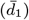 and the determinant 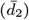, and the permanent 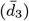 of its diversity matrix *D*.

## 5 Acknowledgments

This research was funded by the European Union’s Horizon 2020 research and innovation programme under the Marie Skłodowska-Curie grant agreement Nº 945371, the European Union Horizon 2020 Research and Innovation Programme under Grant Agreement No. 945539 Human Brain Project(HBP) SGA3 and by UiO:Life Science through the 4MENT convergence environment. Figure 3 has been made with Biorender.com through an academic license.

## 6 Code Availability

The LFDeep: Multitask Learning of Biophysically-Detailed Neuron Models project code is hosted on GitHub. The repository includes the implementation of the paper’s three multitask learning architectures for the desitllation of biophysically-detailed neuron models. To access the latest version of the code and contribute to the project, please visit the official GitHub repository at https://github.com/Jonas-Verhellen/LFDeep.

## 7 Supporting Information

### 7.1 Measuring Diversity in MMoE and MMoEEx

In principle, MTL problems are expected to benefit from higher representational diversity, by which we mean that the representations provided by the different experts capture different aspects of the input data. At least in theory, the MMoEEx architecture was constructed to promote higher diversity among representations, however as mentioned before, we did not observe any clear relationship between diversity and improved performance as measured by the expert diversity metrics discussed in the Methods and Materials section. Moreover, after retraining our models we observed that none of the diversity metrics were consistent between runs, see figure 5. Also, note that all three metrics follow the same pattern within a training session. This result supports the point of view that, at least for the task at hand, a large amount of diversity is not required to find a solution.

**Figure 5:**
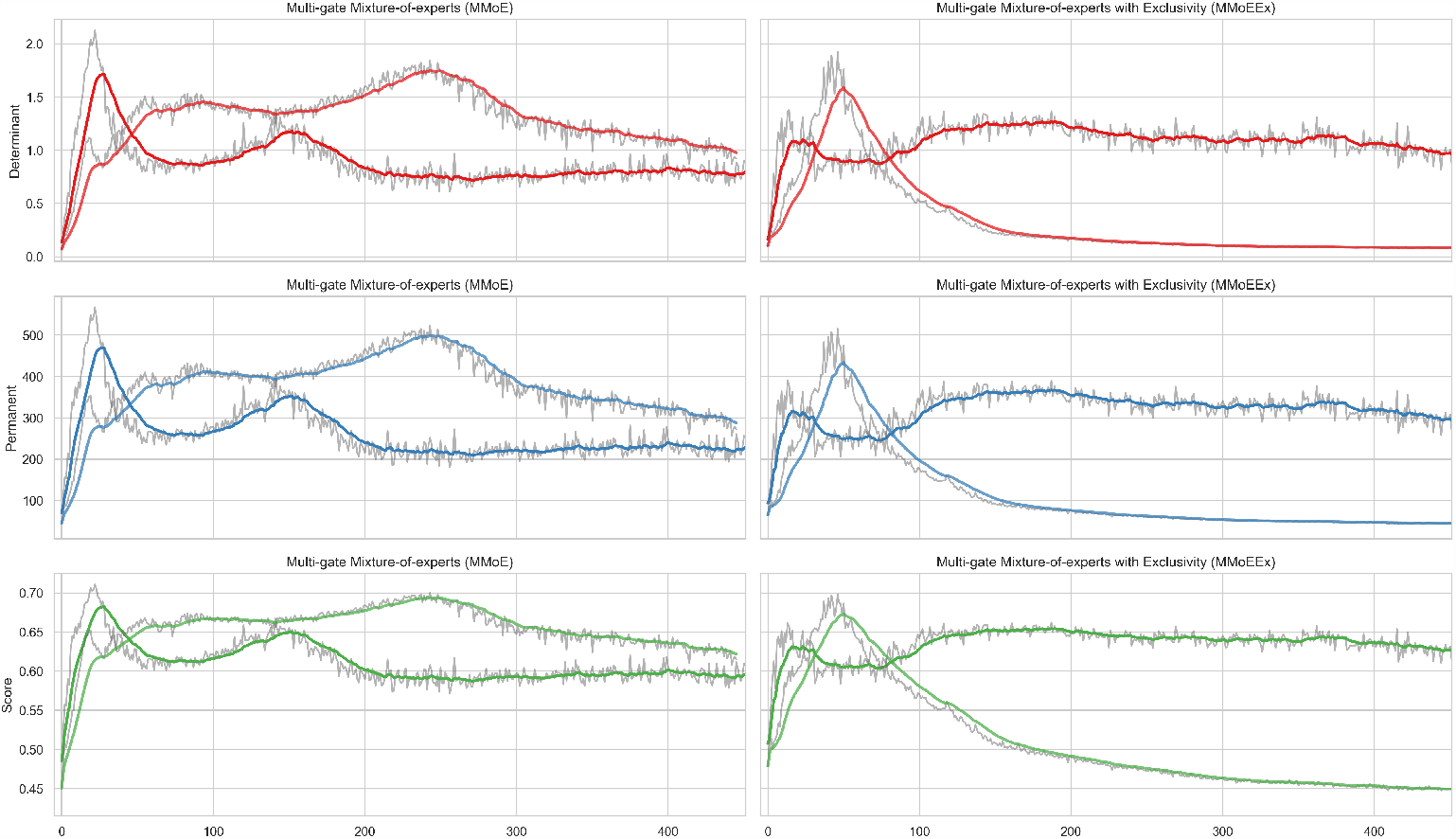
Smoothed time series plot of the evolution of the three diversity metrics (determinant, permanent and score) for two different training runs for both the MMoE (left) and the MMoEEx models (right).

**Figure 6:**
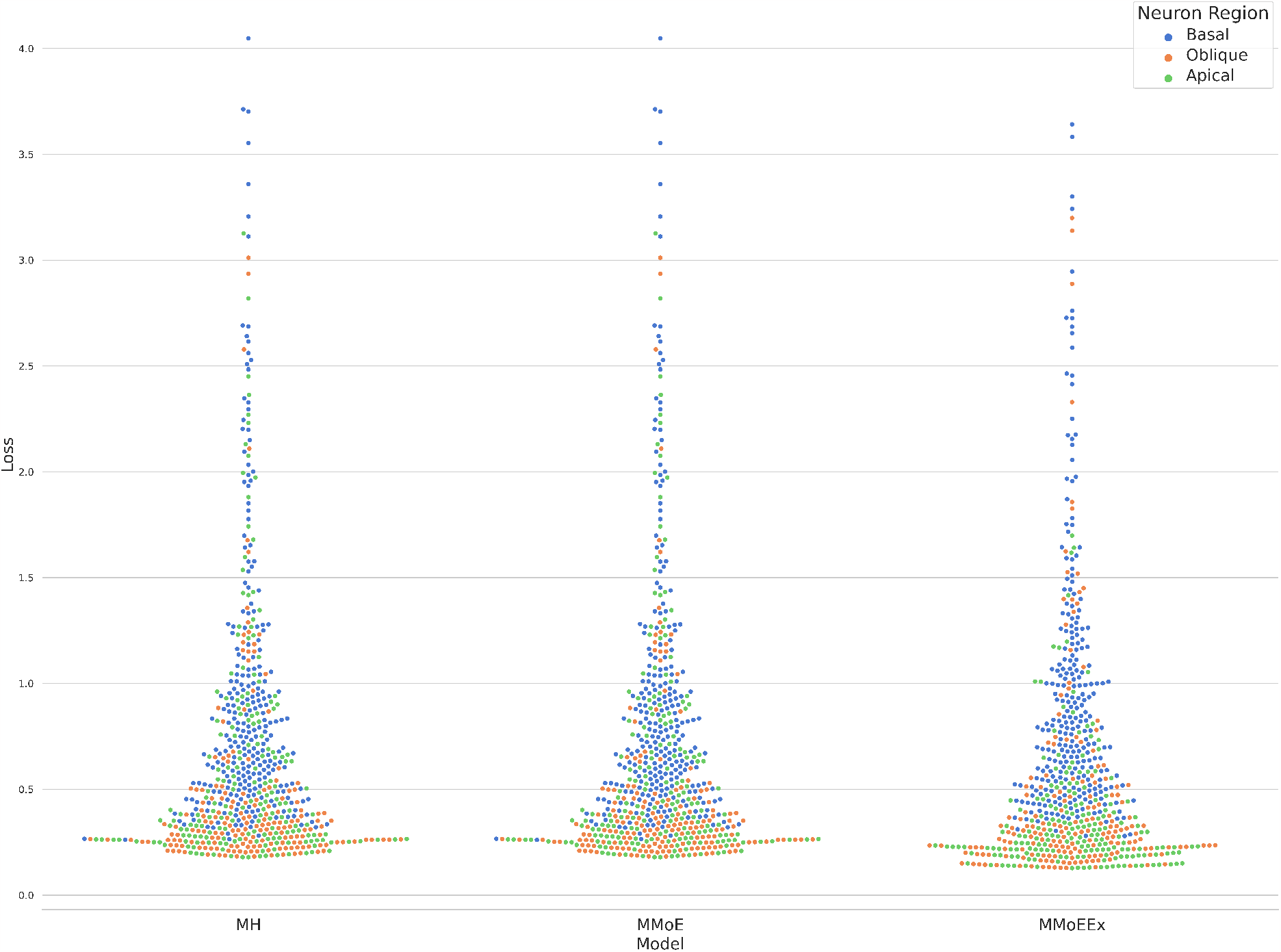
Swarm plots of the generalisation loss for each of the compartments (basal, oblique, apical) of the for each of the three LBTW-trained models: Hard Parameter Sharing (MH), Multi-gate Mixture-of-experts (MMoE), and Multi-gate Mixture-of-Experts with Exclusivity (MMoEEx).

### 7.2 Loss-Balanced Task Weighing for MMoE and MMoEEx

Despite the overall effectiveness of soft parameter models in achieving high average task performance, it is important to acknowledge the potential for negative transfer learning which could lead to a multi-task model underperforming on specific tasks. To address this issue, Loss-Balanced Task Weighting (LBTW) was proposed in the literature as a valuable tool for mitigating potential negative transfer effects. LBTW applies a dynamic update to the task weights in the loss function used during model training. Detailed information regarding the specific loss function and weight updates used by LBTW can be found in the Methods and Materials section. In this appendix, we present results obtained by applying LBTW to the MMoE and MMoEEx architectures, and compare them to the previously described training and generalisation performance of the MMoE and MMoEEx architectures without task-balancing.

The hard parameter model trained with LBTW achieved a minimum validation loss of 0.718, which is higher than the loss of 0.680 obtained through the standard training procedure. Because LBTW was originally designed for soft parameter sharing models only, the lack of improvement in this case was not unexpected. However, the LBTW-trained MMoE and MMoEEx models unfortunately also exhibited higher validation losses compared to the standard training procedures. More specifically, the LBTW-trained MMoE model achieved a validation loss of 0.626, while the standard training procedure resulted in a loss of 0.533. Similarly, the LBTW-trained MMoEEx model had a validation loss of 0.634, whereas the standard training procedure yielded a lower validation loss of 0.551. For a visual representation of the individual validation losses at the end of the training run, refer to Figure 1 of this appendix.

### 7.3 Calculation of Extracellular Potentials

The membrane currents passing through cellular membranes in the vicinity of an electrode give rise to measurable extracellular potentials. One of the downstream applications of the distilled biophysically-detailed models we have presented in this paper is predicting such extracellular potentials, as was shown in Figure 1 of the main text. In this appendix we discuss the calculation of the extracellular potential based on the prediction of our multi-task learned models in more detail. To calculate them we rely on volume conductor theory [49], which dictates that assuming an infinite homogeneous and isotropic extracellular medium, the potential *ϕ* resulting from a multi-compartment neuron model is given by,

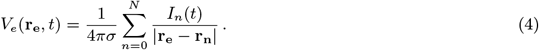

In this equation [46], *I*_*n*_(*t*) denotes the transmembrane currents in a compartment positioned at **r**_**n**_, and **r**_**e**_ indicates the the position of the electrode. The sum runs over all N compartments of the biophysically-detailed neuron model, and *σ* is the extracellular conductivity which is experimentally determined to be around 0.3*Sm*^*−*1^ for cortical grey matter [12]. The multi-task learning models predicted membrane potentials, and not membrane currents, but given full knowledge of both the membrane potentials and the cell model itself, the corresponding membrane currents could be directly inferred (see the LFPy documentation regarding the method “get_transformation_matrix_vmem_to_imem”). Using the inferred membrane currents and the software package LFPy [17], we calculated the resulting extracellular potentials.

Extracellular potentials are a compound result of the contributions of different neuron compartments, and due to the fact that we do not explicitly train our models to predict them, accurate prediction of extracellular potentials is not necessarily a given. While on average our predictions are about as accurate as the experimental variance of the Hay model, some compartments or specific patterns of neural activity may be more or less accurately predicted. For instance as shown in Figure 7 of this appendix, the predictions of membrane voltages in apical and oblique compartments match the ground truth closely, whereas predictions for basal compartments are generally less accurate. As a result of this spatial variance, extracellular potentials near the soma are less accurately predicted in this specific case. Therefore, the use of these distilled biophysically-detailed neuron models in downstream prediction of extracellular potential should be handled with care.

**Figure 7:**
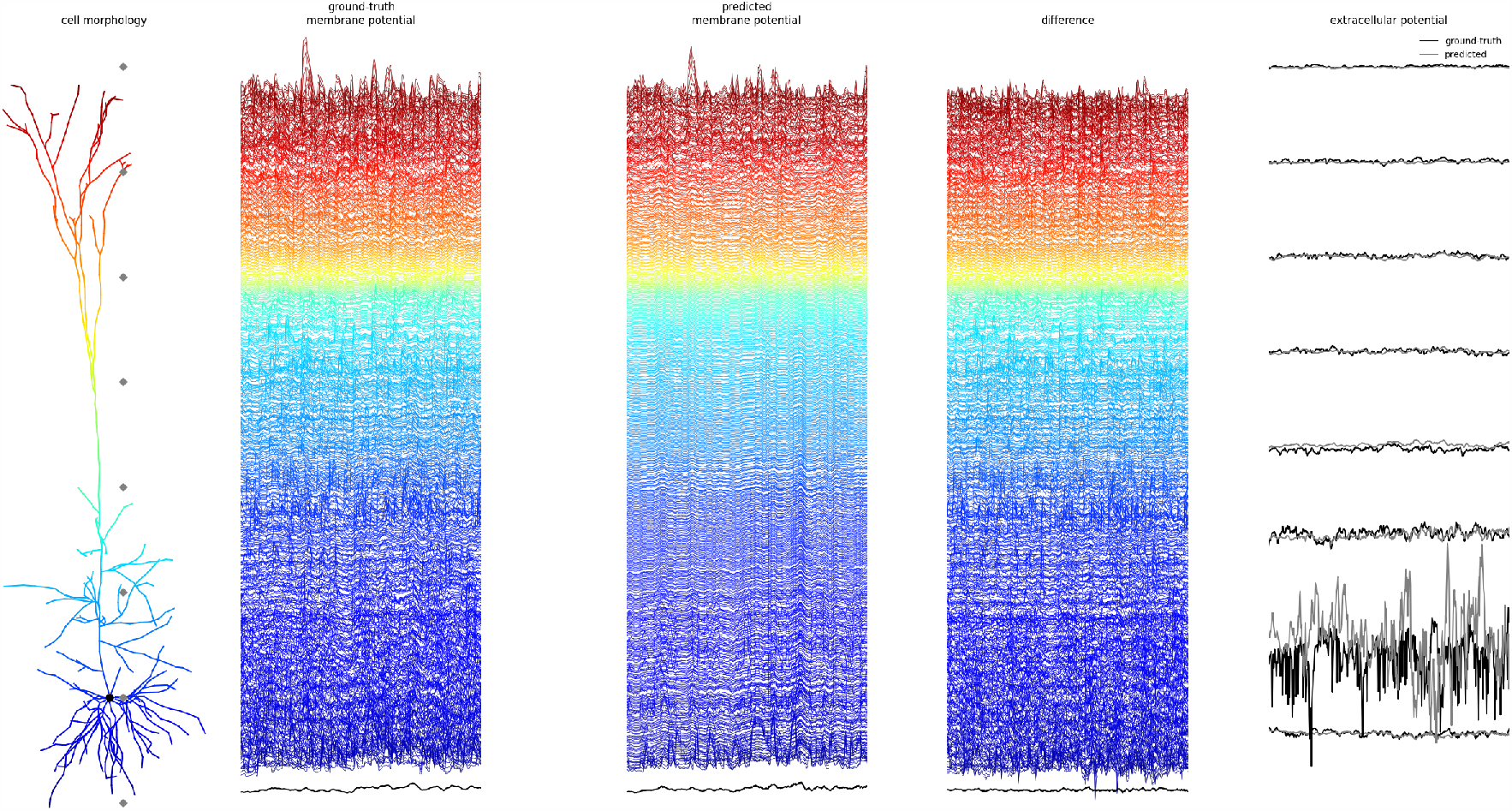
From left to right. Illustration of a biophysically-detailed model of a multi-compartment cortical layer V pyramidal cell model. Membrane voltages as calculated by a biophysically-detailed simulations of the multi-compartment model, used as the ground truth throughout this paper. Membrane voltages as predicted by our best-performing multi-task learning architecture, one time step at a time (1 ms). Comparison between the ground truth and predicted extracellular potentials calculated at eight points representing the position of the electrodes.

